# Structural insights into context-dependent inhibitory mechanisms of chloramphenicol in cells

**DOI:** 10.1101/2023.06.07.544107

**Authors:** Liang Xue, Christian M.T. Spahn, Magdalena Schacherl, Julia Mahamid

## Abstract

Ribosome-targeting antibiotics represent an important class of antimicrobial drugs. Chloramphenicol (Cm) is a well-studied peptidyl transfer center (PTC) binder, and growing evidence suggests it inhibits translation in a nascent peptide sequence-dependent manner. How such inhibition on the molecular scale translates to action on the cellular level remains unclear. Here, we employ cryo-electron tomography to visualize the impact of Cm inside the bacterium *Mycoplasma pneumoniae*. By resolving cellular Cm-bound ribosomes to 3.0 Å, we provide atomic detail on Cm’s coordination and interaction with natural nascent peptides and tRNAs in the PTC. We find that Cm leads to accumulation of translation elongation states that indicate ongoing futile accommodation cycles, and to extensive ribosome collisions. We thus suggest that beyond its inhibition of protein synthesis, the action of Cm may involve activation of cellular stress responses. This work exemplifies how in-cell structural biology advances understanding of mechanisms of action for extensively-studied antibiotics.

## Introduction

Translation is an essential process for every living cell and thus one of the major targets for antimicrobial drugs^1, 2^. Chloramphenicol (Cm) is the first broad-spectrum antibiotic to be clinically used. It inhibits translation through binding to the peptidyl transferase center (PTC) in the large subunit (LSU/50S) of the ribosome^1–3^. Structural studies suggest that Cm blocks new peptide bond formation by sterically hindering positioning of the aminoacyl moiety of the A-site tRNA in the PTC^4–6^. Yet, in contrast to the longstanding notion that Cm is a general inhibitor of translation, recent evidence from ribosome profiling and toeprinting indicates that Cm, and similar PTC-binding antibiotics such as linezolid, block translation in a manner that depends on the amino acid sequence of the nascent peptide^7^. Cm preferentially inhibits peptidyl transfer when the penultimate residue of the nascent peptide (position -1; position 0 defined as the one attached to the P-site tRNA) is alanine (Ala), or to a lower extent, serine (Ser) and threonine (Thr). The presence of aspartic acid (Asp) in position 0 or lysine (Lys) in position -3 potentiate Cm inhibition. Conversely, Cm shows almost no inhibition when glycine (Gly) is in position 0 or in the incoming aminoacyl-tRNA (aa-tRNA; position +1) in the A site^7^. These findings are confirmed by *in vitro* single-molecule fluorescence experiments, showing that Cm does not inhibit translation until the arresting sequence motifs are synthesized, and that inhibition is circumnavigated when the incoming aa-tRNA carries Gly^8^. Single-molecule tracking of translation kinetics in *Escherichia coli* cells treated with Cm also indicate slow but ongoing translation^9^. Thus, despite Cm being one of the most studied antibiotics, how the new evidence on the context-dependent mechanism of action are manifested at a structural level remains an open question.

To address this gap, recent *in vitro* structures of *Thermus thermophilus* ribosomes with non-hydrolyzable tripeptidyl-tRNA analogs as P-site ligands reveal favorable context-dependent interactions between Cm and amino acid residues of the nascent peptide, especially Ala in the position -1, that are required to stabilize Cm in the PTC^10, 11^. A similar sequence-dependent inhibition mechanism is also postulated for oxazolidinone antibiotics that bind at a PTC site overlapping with Cm^12^. However, because fragments mimicking aa-tRNA outcompete Cm from its canonical binding site^10^ and only A-site bound Gly-tRNA is compatible with simultaneous Cm binding^11^, the structure of a Cm-stalled ribosome complex with full peptidyl-tRNA in the P site and aa-tRNA in the A site remains elusive. Furthermore, *in vitro* experiments with defined short peptide mimics cannot recapitulate the inhibition of translation of the plethora of mRNAs inside living cells.

In this study, we used cryo-electron tomography (cryo-ET) to image intact *Mycoplasma pneumoniae* cells treated with Cm, and obtained in-cell maps at better than 3.0 Å resolutions by sub-tomogram analysis. This enabled us to analyze the interaction of Cm with translating ribosomes in high detail, and decipher the impact of Cm on the translation processes across scales within the native cellular context.

## Results

### High-resolution structural features of ribosomes in bacterial cells

Cryo-ET data of *M. pneumoniae* cells treated with Cm for 15 minutes was acquired at a pixel size of 1.33 Å, with each tomogram capturing the majority of one cell (Methods). Subjecting 30,774 ribosome-containing sub-tomograms from 139 cells to structural analysis resulted in a 70S ribosome map at 3.0 Å global resolution, with an overall B factor^13^ of 85 Å^2^ (Extended Data Fig. 1). Focused refinements on the large (50S) and small (30S) ribosomal subunits provided maps at 2.9 Å (with the core resolved to the Nyquist limit) and 3.2 Å overall resolution, respectively (Fig. 1 and Extended Data Fig. 1d-f). The enhanced resolution of the maps, compared to our previous work^14, 15^, allowed us to improve the atomic model for *Mycoplasma* ribosomes (Extended Data Tables 1-3). For instance, we modeled a number of naturally bound polyamine molecules, namely putrescine (PUT), cadaverine (N2P), spermine (SPM) and spermidine (SPD), as well as magnesium and potassium ions (Fig. 1, Extended Data Fig. 1j-l and Extended Data Table 4). Polyamines were shown to bind to ribosomes in *in vitro* studies, and are important for stabilization of the ribosomal RNA (rRNA) structure and translation regulation^16^. Our structures demonstrated the natural composition of polyamines associated to ribosomes inside *M. pneumoniae* cells. While *M. pneumoniae* lost essential enzymes to synthesize polyamines, it preserves membrane transporters responsible for polyamines uptake from the environment or host^17, 18^. We also identified and modeled several rRNA base modifications (Fig. 1 and Extended Data Table 4), of which six align with the modifications identified in cryo-EM and X-ray maps of isolated *E. coli or T. thermophilus* ribosomes^19–22^. We found one base to be differently modified (16S rRNA m^4^C1377 vs. m^4^Cm1402 in *E. coli* and *T. thermophilus*) and one to be unique to *M. pneumoniae* (23S rRNA m^1^G783). In addition, the C-terminal domains (CTDs) of ribosomal proteins S6 (aa131-184) and L31 (aa47-100) were better resolved in the new maps and could be correctly modeled. The proteins L31 and S19 form intersubunit bridge B1c in bacterial ribosomes^23^. Interestingly, the 20 most C-terminal residues of L31 (an extension in *M. pneumoniae* compared to other bacteria^15^) reach even further to contact protein S10 (Fig. 1), thereby strengthening the intersubunit bridge.

**Fig. 1.**
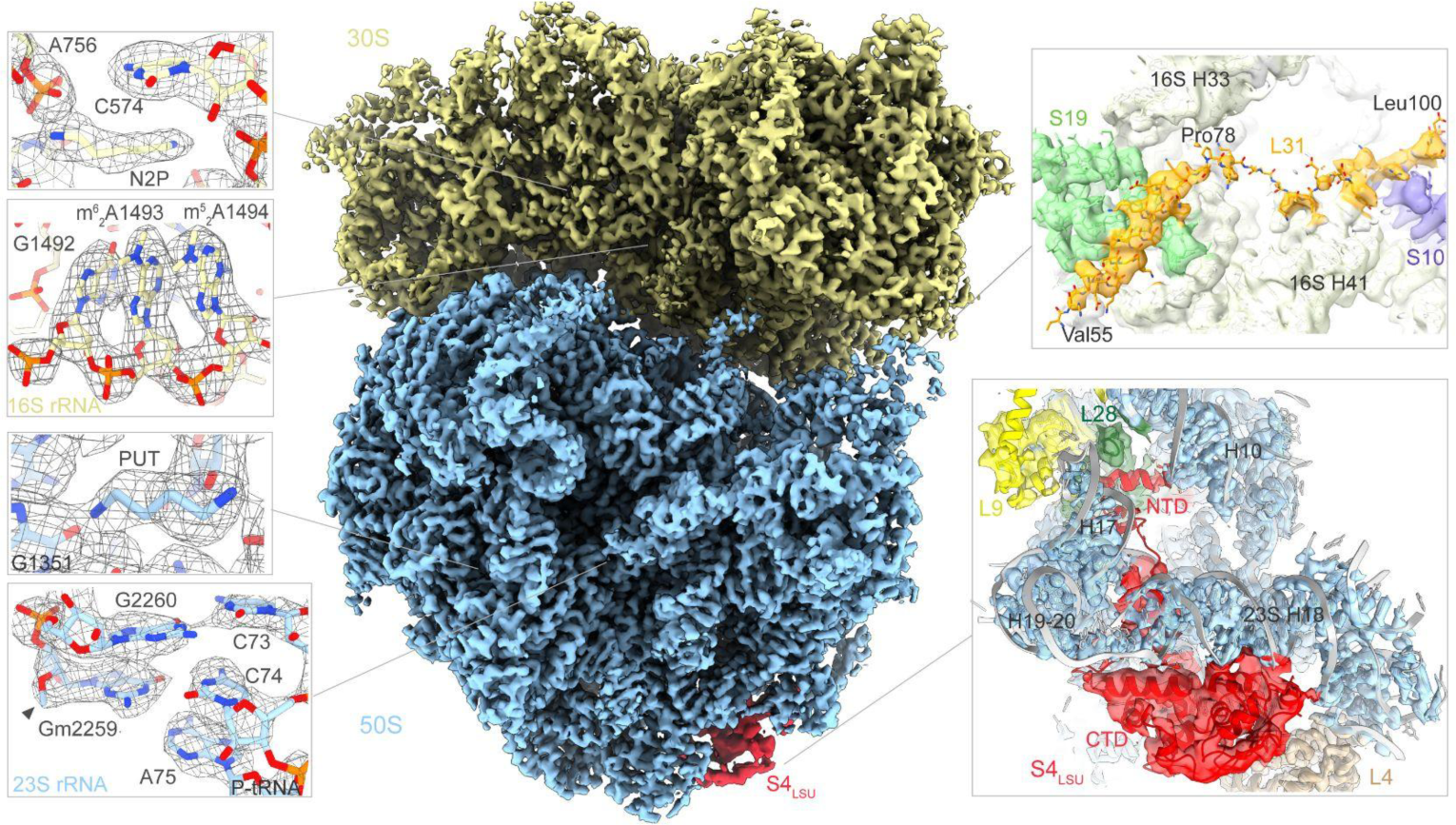
Ribosome maps and models in Cm-treated *M. pneumoniae* cells. Composite of the focused refined 50S and 30S maps of *M. pneumoniae* 70S ribosome. Left insets: representative high-resolution features of the polyamines cadaverine (N2P), putrescine (PUT), and several RNA-base modifications of 16S rRNA (m^6^_2_A: N^6^,N^6^-dimethyladenosine) and 23S rRNA (Gm: 2’-*O*-methylguanosine). Right insets: CTD of the ribosomal protein L31 was modeled from Val55 to Leu100. The interacting rRNA and ribosomal proteins are indicated. A second copy of ribosomal protein S4 was resolved and identified on the large ribosomal subunit (S4_LSU_) in about one third of 70S ribosomes after focused classification. Map and model around S4_LSU_ are shown.

Surprisingly, in approximately one third of all 70S ribosomes, we found a second copy of the small ribosomal protein S4 bound to the large ribosomal subunit (hereafter called S4_LSU_; Fig. 1 and Extended Data Fig. 2a-d). The conformation of this additional S4_LSU_ differs from that of the canonical S4 in the small ribosomal subunit or S4 in the RNA polymerase anti-termination complex^24^ (Extended Data Fig. 2f,g). The S4_LSU_ binding site is located on helices 12, 13 and 18 of 23S rRNA, a site that has not been previously reported as a factor-association site in bacterial ribosomes, and is far away (about 200 Å) from S4’s canonical binding site near the mRNA entry channel. The flexible NTD of S4_LSU_ (aa1-30) protrudes into a cavity located below 23S rRNA helix 18 and contacts protein L28 (Fig. 1 and Extended Data Fig. 2e). The outer perimeter of the cavity is formed by 23S rRNA helix 15, which is similar to that in *Bacillus subtilis* in length and conformation but is missing in *E. coli* (Extended Data Fig. 2h-k). Ribosomes with the S4_LSU_ showed no obvious difference in their distribution across translational states, polysome association, or subcellular regions, compared to the overall 70S population (data not shown). S4_LSU_ was also found in both 70S and free 50S after focused classification of the untreated *M. pneumoniae* ribosomes from our previous study^15^, with the same occurrence frequency and overall structure (Extended Data Fig. 2b,c). S4_LSU_ thus appears to naturally occur in *M. pneumoniae,* but not directly involved in the translation process.

**Fig. 2.**
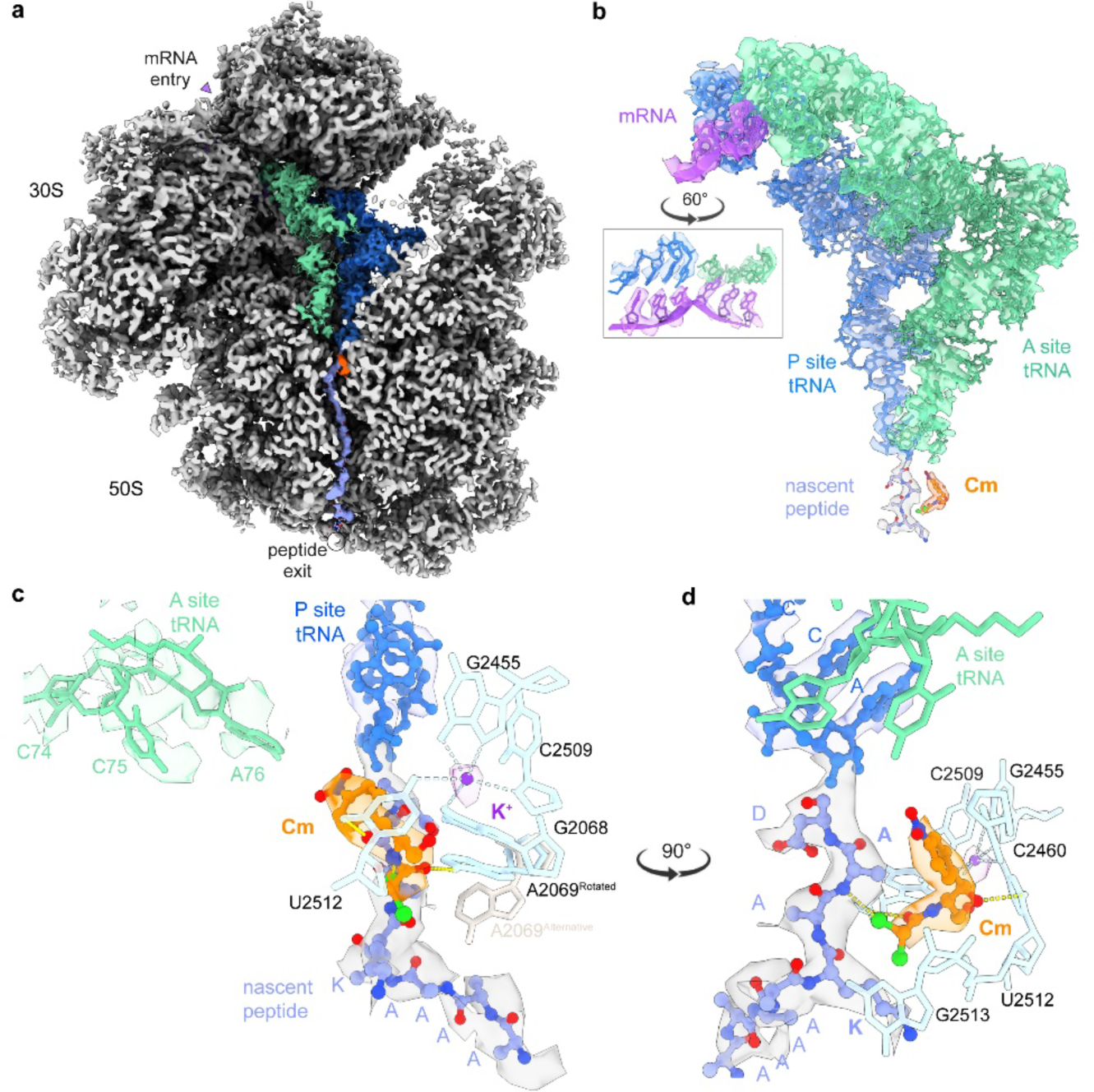
The Cm binding site in PTC is shaped by 23S rRNA, ions and the nascent peptide. **A,** In-cell ribosome map resolves mRNA, tRNAs and the native nascent peptide. Differently sharpened and blurred maps were combined to visualize the nascent peptide density (blue grey) from the PTC to the peptide exit site. **B,** Atomic model for mRNA (purple), A- and P-site tRNAs (green and blue, respectively), nascent peptide (blue grey) and Cm (orange). Inset: condon-antidon pairing in the decoding center. **c-d,** Cm binding pocket in the A site of the PTC is formed by 23S rRNA nucleotides (cyan; only a few bases are displayed) and the nascent peptide (blue grey). For clarity, only positions 0 to -7 of the nascent peptide are shown. The first four residues (DAAK) were modeled in accordance to a ribosome profiling study^7^. A density (purple grey) near the Cm binding site was resolved and was modeled as a K^+^ ion based on previous studies^25, 26^. The CCA-tail of aa-tRNA in the A site is shown with the corresponding density.

### Cm interacts with the native nascent peptide in the PTC

The in-cell consensus map resolved to the data limit of 2.7 Å at the 50S core enabled us to investigate Cm’s mechanism of action in atomic detail in the context of stalled native translation complexes. Cm was clearly resolved in the A site of the PTC (Fig. 2a), its canonical binding site consistent with previous *in vitro* structures^4–6, 10, 11^. Ribosomal ligands, including mRNA, aa-tRNA in the A site and the natural peptidyl-tRNA in the P site (in contrast to the synthetic peptide analogs required to generate *in vitro* structures^10, 11^) could be modeled with high confidence (Fig. 2b). We found the body of aa-tRNA to be fully accommodated, except for its CCA-tail carrying the incoming amino acid which was positioned about 10 Å away from the A site cleft due to Cm’s steric hindrance (Fig. 2c). The blurred local density indicated that the CCA-tail is not stably positioned and may adopt different conformations (Extended Data Fig. 3a). Connecting to the CCA-tail of the P-site tRNA, the nascent peptide could be traced from the PTC to the nascent peptide exit site, with side chain densities resolved for amino acid residues from position 0 to position -3 (Fig. 2a,c). Hence, amino acid residues at positions 0, -1 and -3 were built as Asp, Ala, and Lys, in line with the sequence reported to be overrepresented in Cm-stalled ribosomes^7^. The remaining nascent peptide after position -4 could be traced only by backbone and was modeled as poly-Ala (labeled as poly-UNK in the model).

**Fig. 3.**
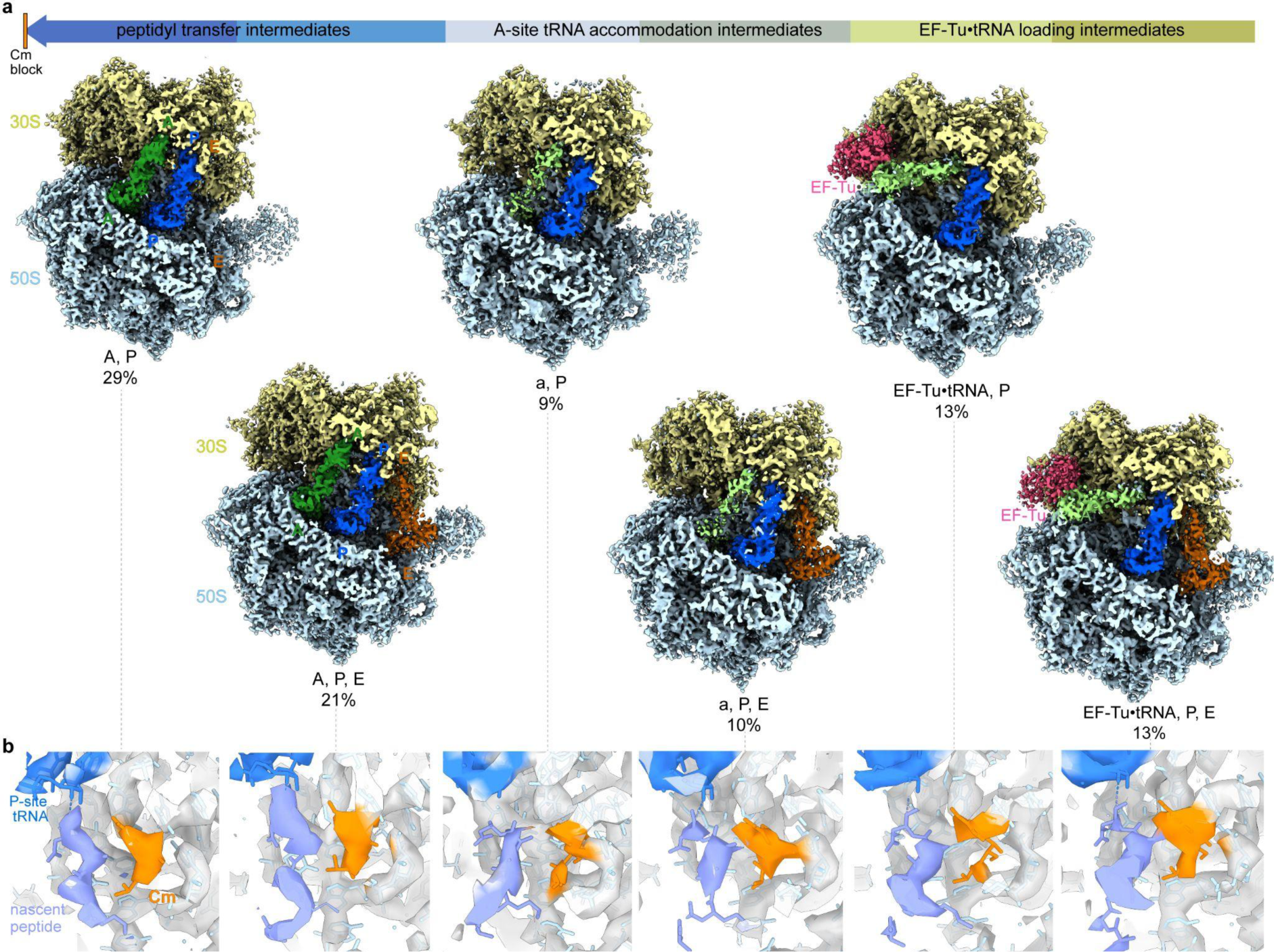
Enriched translation intermediates prior to peptidyl transfer in Cm-treated cells. **a,** Seven translation elongation intermediates (six of these are shown) are characterized by differences in tRNAs (light green: A/T site; dark green: A site; blue: P site; brown: E site) and the elongation factor EF-Tu (red) binding (Extended Data Fig. 4). A unique class name, for example ‘A, P, E’, was given to each class. The percentage of each class was calculated based on particle numbers. The lower case ‘a’ refers to the flexible aa-tRNA near the A site, which is only partially resolved (detailed in Extended Data Fig. 4g). The seventh rotated pre-translocational class ‘A*, P/E’ is not shown here (detailed in Extended Data Fig. 4i). **b,** Cm (orange for density and fitted molecule) was resolved in all six major classes, with the corresponding density observed in the canonical binding site in PTC after fitting the model.

In the PTC, 23S rRNA nucleotides (A2459, C2460, G2068, A2069, U2512) form the binding pocket for Cm. The base of C2460 and Cm’s nitrobenzene ring interacts via π-stacking (Fig. 2d). The base of A2069 (equivalent to A2062 in *T. thermophilus* and *E. coli*) rotates by more than 120° upon Cm binding when compared to the untreated *M. pneumoniae* ribosome structure^15^ (Extended Data Fig. 3c. d), consistent with previous observations^4–6^. In the rotated conformation, A2069 forms hydrogen bonds with Cm’s carboxyl group and the main chain N atom of the residue at position -2 (Fig. 1c, d). We also observed additional density corresponding to other possible rotamers of A2069 (Extended Data Fig. 3c,f-j), reflecting its dynamic nature.

In addition to 23S rRNA, we found that the elongating nascent peptide directly interacts with Cm. The side chain of position -1 residue forms a CH–π interaction with the nitrophenyl ring of Cm, and the residue’s main chain N atom also forms a hydrogen bond with the CL1 atom of Cm (Fig. 2d). This model supports a central role of the position -1 residue in the interaction with Cm, consistent with previous *in vitro* structures^10^. Moreover, we found that the residue at position -3 is involved in shaping the Cm binding pocket in the native translation complex (Fig. 2c,d), restricting the pocket at the side facing the nascent peptide tunnel. We modeled the nascent chain position -3 residue as Lys, consistent with the previous functional study^7^ which revealed a more pronounced translation inhibition by Cm upon accumulation of Ala at position -1 and Lys at position -3. The lysyl side chain is located at a distance of about 3.5 Å from Cm’s dichloroacetyl group and can be stabilized by aliphatic-aromatic stacking on 23S rRNA nucleotide G2513. From position -4 onwards, the nascent peptide is kinked by nearly 90 degrees, which places it far away from the PTC and the Cm binding site (Fig. 2c and Extended Data Fig. 3c-j).

In *T. thermophilus*, Cm was postulated to interact via its (methylene) C4-hydroxyl group with a potassium ion in the PTC (2.7 Å distance between Cm’s O4 and K^+^; PDB 4V7W^4^). We were able to place a corresponding K^+^ ion, coordinated by 23S rRNA nucleotides U2512, G2068, G2455 and C2509, 4.1Å away from Cm’s methylene hydroxyl (O4::K^+^ distance; Fig. 2c,d). This distance does not allow for direct ion coordination, but implies an indirect interaction via a water molecule, in agreement with other Cm-bound *T. thermophilus* ribosome structures (4.1 Å for PDB 6ND5^6^; 4.25 Å for PDB 7U2J^11^). In summary, our in-cell structural model provides detailed information about how Cm binds and reshapes the PTC, and its interactions with the natural nascent peptide that give rise to the sequence-dependent inhibition.

### Cm binding leads to multiple translation elongation intermediates

To assess the general impact of Cm on the translation process, we performed structural classification of the 70S ribosomes in Cm-treated cells. We identified seven significantly populated translation elongation intermediates, among which six were determined at better than 4.5 Å resolution, allowing unambiguous assessment of the presence of the Cm molecule in each of the states (Fig. 3 and Extended Data Fig. 4a-f). We found about 50% of all 70S ribosomes to be stalled in the classical pre-translocational states ‘A, P’ and ‘A, P, E’ (named according to tRNA occupancy; Fig. 3 and Extended Data Fig. 4a-f). As in the consensus structure, the A-site tRNA was found to be fully accommodated except for its CCA-tail, due to Cm’s steric clash with the incoming aa-tRNA. Accordingly, peptidyl-transfer cannot occur, which is expected due to the inhibited peptide bond formation^1, 2^. In addition, 19% of 70S showed a weak tRNA density near the A site (classes ‘a, P’ and ‘a, P, E’) in which only the tRNA anticodon loop bound to mRNA in the 30S decoding center showed clear density, whereas the tRNA main body density was blurred (Fig. 3 and Extended Data Fig. 4g). The blurred local density suggests that the incoming aa-tRNA possibly swings between the A/T and A sites (Extended Data Fig. 4h). Two additional decoding intermediates with bound EF-Tu•tRNA (classes ‘EF-Tu•tRNA, P’ and ‘EF-Tu•tRNA, P, E’) accounted for 26% of the ribosomes, similar to the fraction classified in untreated cells^15^. Blurred local density for EF-Tu suggests that the respective subpopulation is a mixture of decoding intermediates, which could not be further classified. The above described six intermediates can be aligned along the translation elongation trajectory prior to new peptide bond formation (Fig. 3a). Remarkably, they all showed a clear Cm density in their PTCs (Fig. 3b).

**Fig. 4.**
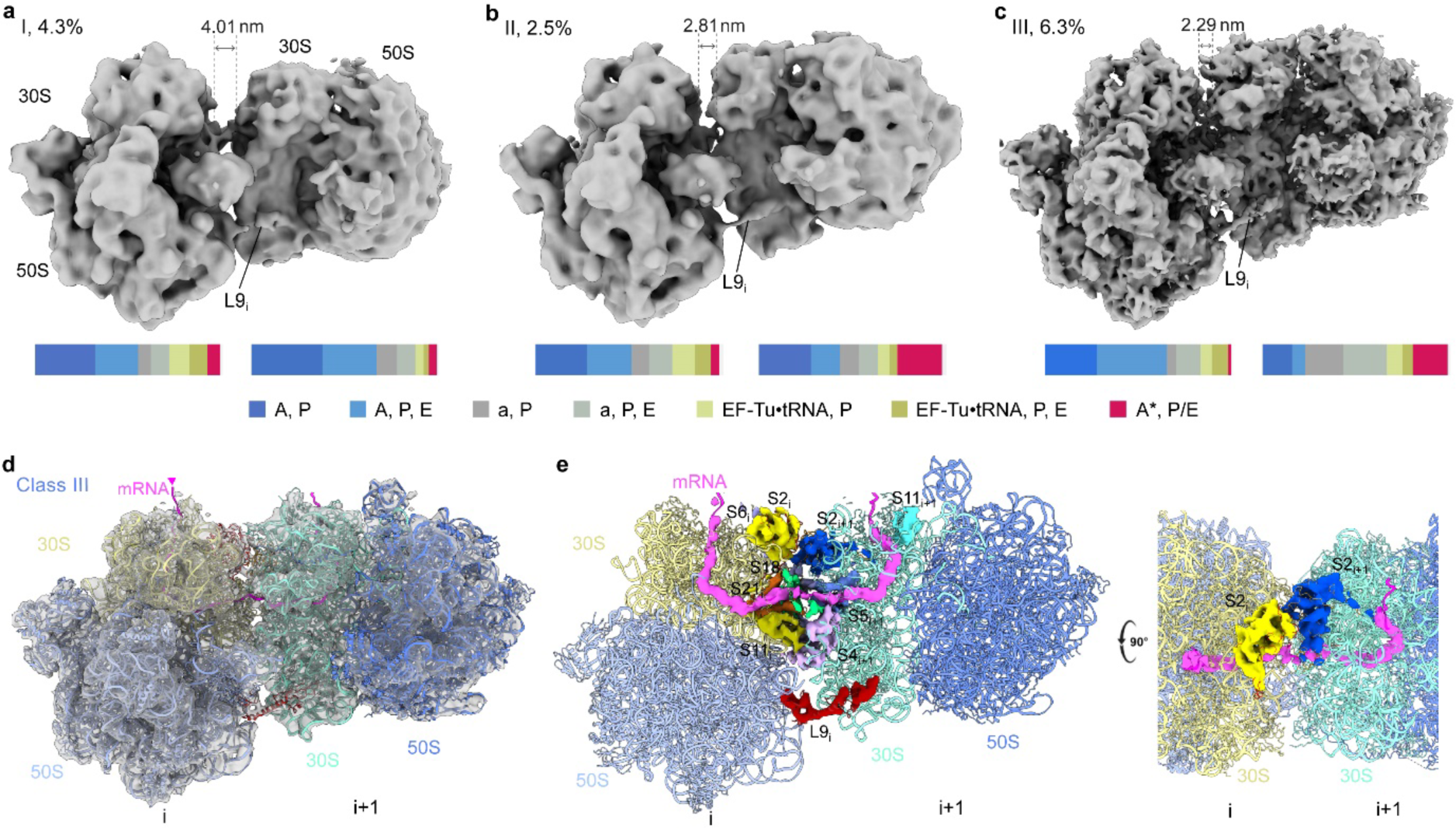
Polysome arrangements in Cm-treated cells. Three different polysome pair (disome) arrangements found in *M. pneumoniae* cells treated with Cm: disome classes I (**a**), II (**b**) and III (**c**), each shown with its percentage among all 70S ribosomes and mRNA-exit-to-entry distance (dashed lines; see details in Methods). Lower bars: distributions of the translation elongation states for the leading and the following ribosomes of each disome class are calculated based on the ribosome state classification results (Extended Data Fig. 4a). **d-e**, Map and atomic model of disome class III. The major ribosomal proteins at the interface are S2, S6, S18, S21, S11 and L9 of the leading ribosome (i), and S2, S5 and S4 of the following ribosome (i+1). Ribosomal protein S1 is not found in *M. pneumoniae*^17^. The mRNA (pink) path can be traced threading between ribosomal proteins S21/S18 of the leading ribosome and S4/S5 of the following one.

In addition, we identified a rotated pre-translocational intermediate (’A*, P/E’) with a marginally translocated A-site tRNA and a hybrid P/E-site tRNA, accounting for 2.6% of the 70S ribosomes refined to 7.6 Å resolution (Extended Data Fig. 4i). The fragmented density for tRNA in the A-site indicates that this subpopulation is a mixture of rotated-1 and -2 pre-translocational states (also called hybrid states H2* and H1, respectively)^27^. While we could not determine whether Cm is bound in the PTC due to the low resolution of this map, formation of the rotated pre-translocational states with hybrid P/E-site tRNA clearly require successful peptidyl transfer. This intermediate could therefore represent the fraction of ribosomes where Cm did not inhibit peptide bond formation due to the presence of a contextually disfavored residue, like Gly at position 0 of the nascent peptide^7^. In summary, inside living cells, Cm binding to ribosomes enriches sequential translation elongation intermediates prior to new peptide bond formation.

### Cm’s action is associated with polysome reorganization

To probe how Cm influences translation at the cellular level, we performed spatial analysis of the ribosome classes mapped back into the 3D cellular volumes, and particularly assessed their arrangement in polysomes. In comparison to untreated cells where 26.2% of 70S ribosomes were found to associate in closely-assembled polysomes^15^, only 15.7% of ribosomes were annotated as polysomes in the Cm-treated cells (Extended Data Fig. 5a-e). 91.3% of the ribosome pairs (disomes) within the polysomes exhibited the compact “top-top” (“t-t”) configuration^15, 28^ (Fig. 4). Disome sub-tomograms were extracted and subjected to classification, resulting in the determination of three distinct arrangements (class I, II and III; Fig. 4a-c and Extended Data Fig. 5f-h). The three classes differed in the relative rotation and displacement of the following ribosome with respect to the leading one, which can be reflected by the distance from the leader’s mRNA exit to the follower’s mRNA entry site (Extended Data Fig. 5i).

In all the three disome classes found in Cm-treated cells, ribosomal protein L9 of the leading ribosome adopts an extended conformation that reaches out to the following ribosome (Extended Data Fig. 5f-h), similar to that found in the “t-t” disomes in untreated cells^15^. Disome class I is structurally similar to the “t-t” disomes in untreated *M. pneumoniae* cells, while classes II and III are more compact (Fig. 4a-c and Extended Data Fig. 6a-c). The most compacted disome class III was resolved to 8.7 Å, within which mRNA could be traced and modeled from the leading to the following ribosome, as well as the ribosomal proteins within the interface (Fig. 4d-e). This class III resembles the recently reported structures of *in vitro* collided ribosomes of both *B. subtilis*^29^ and *E. coli*^30^, in terms of the overall structure, mRNA trajectory and interface proteins (Extended Data Fig. 6d-g). Our disome maps, however, do not contain additional density that could be assigned to the rescue factor SmrB or MutS2 found in the *in vitro* collided disomes^29, 30^, in line with *M. pneumoniae’*s genome lacking SmrB or other small MutS-related (SMR) domain containing proteins^17^. Other minor differences at the disome interface originate from *M. pneumoniae’s* specific set of amino acid extensions in proteins S5, S6 and S18, as well as the lack of protein S1^15, 17^.

Integrating the polysome analysis with the results from our classification of ribosome translation elongation intermediates showed the distribution of states for the leading and following ribosomes to be related to the different disome configurations (Fig. 4). Specifically, there was an enrichment of the rotated pre-translocational ‘A*, P/E’ state in the following ribosomes of the more compacted disome classes II and III. In contrast, the state distribution of either the leading or the following ribosomes in the less compact class I showed the same patterns as in mono-ribosomes (Extended Data Fig. 5e). While the resolutions of disome maps were not sufficient to directly determine whether Cm is bound, considering the saturating concentrations of Cm applied (Methods) and the unambiguous identification of the Cm density in the vast majority of translation elongation intermediates (Fig. 3b), with the exception of the ‘A*, P/E’ class, we suggest that Cm is also bound in the majority of polysomes. To corroborate this, disome classes II and III only exist in cells upon Cm treatment, and not in untreated cells. These results indicated that Cm induces ribosome collision within polysomes and changes their functional organization in cells. Ribosome collisions could arises from a scenario wherein the leading ribosome is stalled by Cm on the arresting motifs but the following ribosome is still able to elongate. This is supported by the observation that the rotated pre-translocational intermediate ‘A*, P/E’ post peptidyl transfer is over-represented in the following ribosomes of the two most compacted disome classes II and III, while it distributes evenly in the disome class I that is similar in arrangement to polysomes found in untreated cells.

## Discussion

In this work, we demonstrate that it is feasible to obtain maps with local resolutions better than 3 Å and thus atomic-level detail for large macromolecular complexes within intact cells. The use of a smaller pixel size and the acquisition of a larger dataset in this study compared to our previous reports^14, 15^ enabled us to improve the resolution of the ribosome consensus map, while the overall B factor of 85 Å^2^ remains similar. Our refined in-cell *M. pneumoniae* ribosome structure aligns with high-resolution *in vitro* structures from other bacteria^6, 19, 21, 26, 31^, down to the level of small cofactors and ions.

Yet the finding of S4_LSU_ highlights the existence of different ribosome isoforms inside cells and underlines the importance of obtaining such structures. Besides its canonical role in forming the mRNA entry site on the 30S subunit, S4 is known to play essential roles in suppressing translation initiation of mRNAs from its own α operon^32^, in the transcription anti-termination complex during ribosome biogenesis^24, 33^, and in guiding early 16S rRNA folding and 30S subunit assembly^34, 35^. However, there has been no previous report on the involvement of S4 protein in 50S assembly. In light of the diverse roles of S4 in regulating RNA folding, we hypothesize that S4 can serve as a chaperone in 50S folding and assembly. This action could be specific in mycoplasmas, and S4 may later dissociate over time independent of translation. Alternatively, the additional association site on the large ribosomal subunit can serve as a buffering pool to regulate the available S4 proteins in cells. The S4_LSU_ binding site appears to be structurally conserved in bacterial ribosomes and it may therefore serve common, yet unknown, functions across bacteria. Overall, this result demonstrates the unique potential of cryo-ET in the discovery of moonlighting factors on large complexes inside cells.

Our in-cell structure reveals atomic details of how Cm interacts with the natural constituents of the ribosome PTC and sheds light on the molecular inhibition mechanism of Cm. In addition to the 23S rRNA nucleotides forming the main Cm binding pocket, the structure supports the previously reported importance of a potassium ion coordinated by the pocket-forming nucleotides G2068 and U2512 (Fig. 2c,d). Dependence on K^+^ for Cm binding was first shown biochemically^36, 37^ and then confirmed structurally^4, 25^. Cm binding displaces two or more water molecules, which in untreated ribosomes form the hydrogen bond network with the K^+^ and pivotal nucleotides in PTC^25, 26^. This explains resistance to Cm arising from rRNA mutations, *e.g.* at position G2455 (equivalent to G2447 in *E. coli*), which do not directly interact with Cm but coordinate the K^+^ ion^38–41^, or modification of Cm’s C3 hydroxyl group that is part of the K^+^ coordination network^4, 42^.

Cm is long known to sterically prevent proper positioning of the aminoacyl moiety of the incoming aa-tRNA in the A site of the PTC, leaving the nascent peptide connected to the P-site tRNA (Fig. 2). Our in-cell structure demonstrates that direct interaction between Cm and the nascent peptide, especially its position -1 residue, further stabilizes Cm’s occupation of the A site, consistent with previous *in vitro* structures obtained with synthetic nonhydrolyzable peptide analogs^10, 11^. Large side chains of this penultimate residue can clash with Cm, explaining why Ala and to a lower extent Ser and Thr are over-represented in Cm-bound ribosomes^7^. Additionally, we demonstrate that the position -3 residue is involved in forming the Cm pocket (Fig. 2d), which favors Lys to help seal the binding pocket and stabilize Cm binding^7^. Although Cm can bind to vacant ribosomes^4–6^, inhibition is not effective until the arresting nascent peptide motifs are synthesized to further stabilize Cm’s binding and enhance its blocking activity^7, 8^. In its rotated conformation, 23S rRNA nucleotide A2069 (A2062 in *T. thermophilus* and *E. coli*) interacts with both Cm and the nascent peptide (Fig. 2), and plays an important role in sensing the nascent peptide and stabilizing Cm’s inhibition^6, 10, 43, 44^. In untreated *M. pneumoniae* cells with active translation elongation, A2069 in the average map adopts the unrotated conformation^15^. Thus, Cm’s action goes beyond simple blocking of access of the aminoacyl moiety to the PTC; it reprograms the PTC via imposing changes to rRNA conformation, ion coordination network, and the interaction with specific sequences of the elongating nascent peptide.

As a consequence of the impaired peptide bond formation, Cm also reshapes the functional landscape of ribosomes. We found more than 97% of cellular 70S ribosomes to be present in six different translation elongation states prior to peptidyl transfer, including EF-Tu•tRNA decoding and aa-tRNA accommodation intermediates (Fig. 3). The relative abundance for these states differs markedly from their distribution in native untreated cells^15^. Ribosomes with all three tRNAs bound (A, P and E-sites) in the classical pre-translocational state prior to peptide bond formation were not detected in untreated cells, but account for 21% of all 70S in Cm-treated cells. This indicates a functional link between disassociation of the E-site tRNA and successful peptidyl transfer^45^. These findings further suggest that Cm’s inhibition of peptidyl transfer possibly results in repeated rounds of nonproductive accommodation and dissociation of aa-tRNA in the A site, in agreement with a previous single-molecule FRET study^8^. Similarly, hygromycin A binding in a PTC region partially overlapping with Cm’s binding site, leads to oscillation of the incoming aa-tRNA between the A/T-like and the accommodated positions^46^. Our data provide direct structural evidence for the occurrence of these intermediates upon antibiotic treatment in the native cellular context (Fig. 3). When peptide bond formation is inhibited or slowed down, increased dissociation of aa-tRNA can occur via the route used for kinetic proofreading^47^. Accordingly, futile rounds of ternary complex formation and GTP hydrolysis on the ribosome can occur^48^. These unproductive cycles may contribute to diminishing the intracellular GTP pool. Cm and most ribosome-targeting antibiotics are bacteriostatic drugs that halt cell growth but do not kill bacteria. It was recently reported that *B. subtilis* employs (p)ppGpp-mediated cellular stress response to protect against Cm by lowering the intracellular GTP level, and that increasing intracellular GTP levels enhances Cm lethality^49^. Therefore, on top of protein synthesis inhibition, Cm can turn ribosomes into nonproductive machines that consume energy, possibly contributing to an additional effect of the antibiotic on cellular physiology. It is possible that cells in turn adapt to such additional stress, relying on (p)ppGpp-mediated pathways to decrease the GTP level and suppress the futile accommodation cycles. A combination of PTC-targeting antibiotics with drugs suppressing protective bacterial cellular stress responses may represent a promising direction for future antibacterial treatment development.

Cm treatment further profoundly reorganizes the translation machinery in cells, with particular impact on polysome arrangements. We found that about 70% of the detected polysomes in Cm-treated cells represent disome classes that resemble *in vitro* collided disome structures^29, 30^, but that do not exist in untreated cells^15^. Such collisions increase translation errors such as frameshifting^50, 51^, which can be reduced by the ribosomal protein L9^51, 52^. In native untreated cells, L9 of the leading ribosome interferes with elongation factor binding, especially EF-G, to the following ribosome and thereby mediates polysome coordination during active translation^15^. However, for polysomes in Cm-treated cells, where the leading ribosome is prolongedly stalled on arresting sequences, the following ribosome may be able to complete factor-independent translocation albeit at lower efficiency^53, 54^, and collide with the leading one. Indeed, it has been reported that Cm increases translation errors, especially frameshifting and nonsense suppression but not misincorporation^55^. Hence, we suggest that Cm treatment induces ribosome collisions that contribute to Cm’s effect on the cellular level. As a general sensor and inducer of cellular stress responses, ribosome collision caused by Cm could exert a more detrimental effect on cells than pure inhibition of protein synthesis^56–5^. Cells, on the other hand, may alleviate the effect of Cm through the cellular stress response and ribosome collision rescue mechanisms^56, 59^. Indeed, mutations of the ribosome rescue genes in *B. subtilis*^29^ or *E. coli*^30^ cause increased sensitivity to ribosome-targeting antibiotics including erythromycin and chloramphenicol. Although no ribosome rescue proteins are annotated in *M. pneumoniae,* possibly due to its significant genome reduction, our results suggest a general mechanism for antibiotics that act in a sequence-dependent manner to additionally induce ribosome collisions as part of their cellular mechanisms of action.

To conclude, our study provides a comprehensive understanding of Cm’s effect on the atomic, molecular and cellular levels, and complements the current context-dependent inhibition model of Cm and other PTC-targeting antibiotics. We demonstrate that the context-dependent action of Cm is not only reflected in ribosome stalling on specific nascent peptide sequences, but also propagates to the cellular scale in the native context of polysomes, and possibly associated cellular stress response processes. This work establishes how emerging in-cell structural biology can advance mechanistic understanding of drug action in their natural context – i.e. inside the cell.

## Methods

### Cryo-ET data collection and processing

*Mycoplasma pneumoniae* cell cultivation on grids and sample preparation were performed as previously described^15, 60^. Chloramphenicol (Sigma-Aldrich) was added to the cell culture in the fast-growing phase at 0.2 mg/ml final concentration, and incubated for about 15 minutes before plunge freezing. Cryo-ET data collection was performed on a Titan Krios G3i microscope equipped with a Gatan K3 camera using SerialEM version 3.9^61^. Tilt-series were collected using the dose-symmetric scheme^62^, with the following settings: magnification 64,000x, pixel size on sample 1.329 Å, tilt range -60° to 60° with 3° interval, K3 camera in non-CDS counting mode, target dose rate on camera 20 e^-^ /pixel/second, 10 frames per tilt image, constant exposure time for each tilt and total dose 137 e^-^/Å^2^. For different tilt-series, the target defocus values ranged from 1 to -3.25 μm. In total, 139 tilt-series were used for data processing and analysis.

### Image processing and ribosome structure refinement

Cryo-ET data was processed in Warp v1.0.9^14^, including frame motion correction, CTF estimation, and tilt-series sorting. Tilt-series alignment was done in etomo v4.11^63^ using gold fiducials. Tomograms were first reconstructed at bin 6 (voxel size 7.974 Å) in Warp, after importing the etomo alignments. Ribosome picking was done using template matching in pyTOM v0.9.7.1^64^, and the top 600 or 900 ranking cross correlation hits for every tomogram were extracted depending on the cellular coverage area. In total, 51,783 sub-tomograms were extracted from 139 cellular tomograms at bin 4 (voxel size 5.316 Å) in Warp, and subjected to 3D classification to remove false positives and free 50S in RELION v3.0.8^65^. Finally, 30,774 sub-tomograms of 70S ribosomes were extracted at smaller voxel sizes for the following refinement and classification.

Initial refinement (alignment and averaging) of the 70S ribosome sub-tomograms was performed in RELION v3.0.8^65^. The alignment parameters for all particles and the average map were imported into M v1.0.9 for multi-particle refinement^14^. Structure-based refinement of both geometric (image and volume warping, particle poses, tilt alignment) and contrast transfer function (CTF) parameters was performed for five rounds in M, which also automatically updates the metadata files in the Warp processing folders. After M refinement on the 70S ribosomes with a global mask, focused refinements with 30S and 50S masks were performed simultaneously with only ‘particle pose’ option to improve local map quality. Fourier shell correlation (FSC) calculation between randomly split half subsets, local resolution estimation, and additional postprocessings were performed in M and RELION.

### Atomic model building and refinement of high-resolution ribosome averages

The atomic models of *M. pneumoniae* 30S ribosomal subunit (PDB ID 7OOC) and 50S ribosomal subunit (PDB ID 7OOD)^15^ were initially docked into the density maps with UCSF Chimera^66^ and manually adjusted in Coot^67^. To model regions of different local resolutions, such as intrinsically flexible RNA or the N-terminal parts of the nascent peptide, different B factors in RELION postprocessing^65^ were used to sharpen or blur the density map. As the modelled mRNA represents an average of all native *M. pneumoniae* mRNAs bound to the imaged ribosomes, a random sense mRNA sequence was chosen, except for the three codons, which were adjusted to the anticodons of chosen tRNAs. The tRNAs were selected based on the prevalence of amino acids found in positions +1, 0, and -1 of the nascent chains in *E. coli* ribosomes upon chloramphenicol treatment^7^. Here, position +1 corresponds to the A-site amino acid, position 0 to the P-site/ultimate, and position -1 to the penultimate amino acid of the nascent chain, respectively. A poly-Ala_(<-4)_-Lys_(-3)_-Ala_(-2)_-Ala_(-1)_-Asp_(0)_ polypeptide was modelled as the nascent chain (the poly-Ala part is labeled as poly-UNK in the deposited model, as the identity is unknown). The tRNA sequences and the codon usage were adjusted to match the experimentally determined most frequently used ones in *M. pneumoniae*^68^, namely: deacylated tRNA^Ala^ (cluster trnD; gene: MPNt01; anticodon: UGC, codon: GCC, U•C wobble pair) for the E-site (where present); nascent chain-acylated tRNA^Asp^ (cluster trnB; gene: MPNt11; anticodon: GUC, codon: GAU, G•U wobble pair) for the P-site, and acylated Lys-tRNA^Lys^ (cluster trnE; gene: MPNt28; anticodon: CUU, codon: AAG) for the A-site, respectively. To ensure correct placement and refinement of the acylated CCA-tail of the A-site tRNA, lysinyl-AMP was created (chemical ID AK9) in Coot and restraints were generated using the module ‘eLBOW’^69^ in PHENIX^70^. AK9 was added to the 3’-end of tRNA-Lys, replacing the adenosine at position 76. All models were refined over multiple rounds using the module ‘phenix.real_space_refine’ in PHENIX and interactive model building and refinement in Coot, using libG restraints^71^ for the RNA. The quality of all refined models was assessed using the ‘comprehensive model validation’ function in PHENIX and wwPDB validation server (https://validate.wwpdb.org). The model validation statistics in Extended Data Table 1-3 was calculated using MolProbity^72^. Ribosomal RNA secondary structure representation from the PDB file was done with RNApdbee 2.0 software^73^ and image produced with VARNA^74^.

### Protein identification and atomic model building for ribosomes with S4_LSU_

Ribosomes showing an additional density near 23S rRNA helices 16-18 were sorted out through focused classification with a spherical mask covering the additional density (Extended Data Fig. 2a). Parallel RELION jobs were performed to mitigate variations. After obtaining the density map, we aimed to identify the protein based on its fold. As automated building of alpha-helices and beta-strands into the additional density did not yield a meaningful model, several poly-Ala stretches were placed manually in Coot with the ‘place helix/strand here’ module. Due to the visibility of a few side chains in the density, the directionality of the potential alpha-helices could be deduced. Also some parts of potential beta-strands were built, and all identified polypeptide stretches were connected via random coil stretches to yield a single polypeptide of 123 residues. The model was submitted for a 3D-structure similarity search to the DALI server^75^ against the Protein Data Bank. This search yielded PDB entry 5WNU chain D^76^ as top hit (Z-score 11.2, rmsd 2.4 Å, lali 112), which corresponds to 30S ribosomal protein S4 from *T. thermophilus* (UniProt ID P80373). Based on the search result, we placed a second copy of *M. pneumoniae* 30S ribosomal protein S4 (now called S4_LSU_), N-terminally truncated to residues 47-205, into the additional density, which gave a robust fitting that only required minor adjustment (Extended Data Fig. 2d). The N-terminal 46 residues were built *de novo* into the density in Coot.

### Classification of translation elongation intermediates

3D classification of the 70S ribosomes was performed in RELION v3.0.8 using the sub-tomograms re-extracted in M v1.0.9 after multi-particle refinement. The hierarchical classification strategy and procedure are similar to those described in our previous study^15^, and are illustrated in detail in Extended Data Fig. 4a. At least three tires of RELION classification jobs were performed. First, the 70S ribosomes were classified using a global 70S mask to remove false positives or ‘bad’ particles. Second, ribosomes were sorted according to the different tRNA binding states, using either a focused mask covering the small ribosomal subunit plus all possible translational factor binding regions, or a spherical mask covering the A-, P-, and E-tRNA binding sites. Third, classification was based on the different elongation factor and A/T tRNA binding states, using a spherical mask focusing on that region. 15,332 ribosomes with clear A and P-site tRNAs were first separated, and further classified based on the E-site tRNA density into ‘A, P’ (8,854 particles) and ‘A, P, E’ (6,478 particles). The rest were classified into five major groups: 7,048 ribosomes with EF-Tu•tRNA and P-site tRNA; 5,661 ribosomes with partially resolved A-site tRNA (only the tRNA tip close to the decoding center is well resolved; labeled as ‘a’) and P-site tRNA; 787 with A-site tRNA and hybrid P/E tRNA; 769 ribosomes with less resolved density near the P site and 1,177 ribosomes show dim 30S subunit. The last two classes (with 769 and 1,177 particles respectively) could not result in any interpretable density maps and were thus not further analyzed. The 7,048 ribosomes with EF-Tu•tRNA and P-site tRNA, and the 5,661 ribosomes with partially resolved A-site tRNA, were finally classified based on the E-site tRNA states. In each tier, at least three parallel classification jobs (with same settings, slightly different settings, or with different masks) were performed, and the most stable job was selected to sort the particles. For each particle classification/sorting, follow-up classification runs were performed until no new/different sub-classes emerged.

### Spatial and statistical analysis of polysomes

The polysome annotation procedure was performed as described before^15^, and considers both the relative positions and orientations of ribosomes in the 3D cellular volume. Polysomes in this study only refer to the closely-assembled polysomes with a distance threshold of 7 nm^15^. The distance determined from a manually defined mRNA exit site on one ribosome (the preceding i) to the mRNA entry site of the following ribosome (i+1) was used to determine whether the two ribosomes belong to the same polysome. In total, 4,838 ribosomes were annotated to be within the polysomes.

Additionally, neighboring ribosome-ribosome pairs within the polysomes were sorted based on their relative rotations^15, 28^. 1,959 “top-top” pairs with mRNA exit to entry distance of 3.3 ± 1.5 nm, and 186 “top-bottom” pairs with distance of 5.4 ± 1.1 nm were detected.

Di-ribosomes (ribosome pairs within polysomes) were additionally structurally classified by first extracting the 4,838 annotated ribosomes with a large box (3.1 Å/voxel, box 256) and then applying focused classification with a spherical mask covering the position of the following ribosome. Three disome structures were classified. The functional states of the leading and the following ribosomes within the disomes were mapped by integrating the above ribosome classification results.

### Modeling of compacted disome structure

Modeling of the most compacted disome class III (resolution of the map is 8.7 Å at FSC=0.143) was performed using the built 30S and 50S subunits as the starting models, which were first rigid-body fitted into the density map using Chimera and Coot. Several parts of the models for the leading (i) and following (i+1) ribosomes needed adjustments in Coot to be accommodated in the disome map. For example, residues R121-A137 of protein S6_i_ must fold differently (compared to free 70S) to avoid clashes with protein S5_i+1_. The C-terminus of protein S6_i+1_ detaches from the ribosome and is disordered from residue S168 on, as it would clash with 70S_i_. The C-terminal domain of protein L9_i_ is in an extended conformation and blocks the translation factor binding site of the following 70S_i+1_. The A site of 70S_i+1_ is empty, as the ternary complex cannot form. The A-site finger of the following ribosome was found in two conformations and the monitoring bases (16S_i+1_ rRNA bases 1467-1468) are mainly in the “flipped-in” conformation. We also used information from in-cell cross-linking mass spectrometry^60^ to guide model building where the path of the polypeptide was somewhat unclear, mainly for the flexible termini of some ribosomal proteins. The model for mRNA located in between the two ribosomes was built to accommodate exactly 10 codons (30 nucleotides) between the two A sites, similar to other structures of bacterial collided disomes^29, 30^ and matches biochemical studies^51^, ranging from 8-10 codons.

Structural visualization and figure generation were using Chimera^66^ and ChimeraX^77^. Statistical analysis and plotting was performed in MATLAB 2016b.

#### Reporting summary

Further information on research design is available in the Nature Research Reporting Summary linked to this article.

### Data availability

Detailed information for all maps and models generated in this work is provided in Extended Data Tables 1-3. The raw cryo-ET data is deposited in the Electron Microscopy Public Image Archive (EMPIAR) under accession code EMPIAR-11520. Maps are deposited in the Electron Microscopy Data Bank (EMDB) under accession codes: 17132, 17133, 17134, 17135, 17136, 17137, 17138, 17139, 17140, 17141, 17142, 17143, 17144, 17145, 17146, 17147. Atomic models are deposited in the Protein Data Bank (PDB) under accession codes: 8P6P, 8P8B, 8P7X, 8P7Y, 8P8W, 8P8V. The depositions will be released upon publication. Maps and atomic models used from previous studies were obtained from the PDB (7OOC, 7OOD, 7P6Z, 5WNU, 6QNR, 7N1P). The Mycoplasma pneumoniae M129 protein and RNA sequences were obtained from NCBI Reference Sequence NC_000912.1.

## Acknowledgements

We thank the EMBL cryo-EM platform, Thomas Hoffmann and EMBL IT for technical support, and Nassos Typas and Olivier Duss for valuable input on the manuscript. C.M.T.S. thanks Charité - Universitätsmedizin Berlin and the Deutsche Forschungsgemeinschaft for funding. J.M. acknowledges support from the EMBL and a Chen-Zuckerberg Initiative grant for Visual Proteomics.

## Author contributions

L.X. and J.M. conceived the study; L.X. collected cryo-ET data and performed structural analysis; M.S. built the atomic models and together with C.M.T.S. assisted interpretation; L.X. and J.M. wrote the manuscript with inputs from all authors.

## Additional information

Extended Data is available for this paper.

No supplementary information is included in this manuscript.

Correspondence and requests for materials should be addressed to

## Extended Data

**Extended Data Fig. 1.**
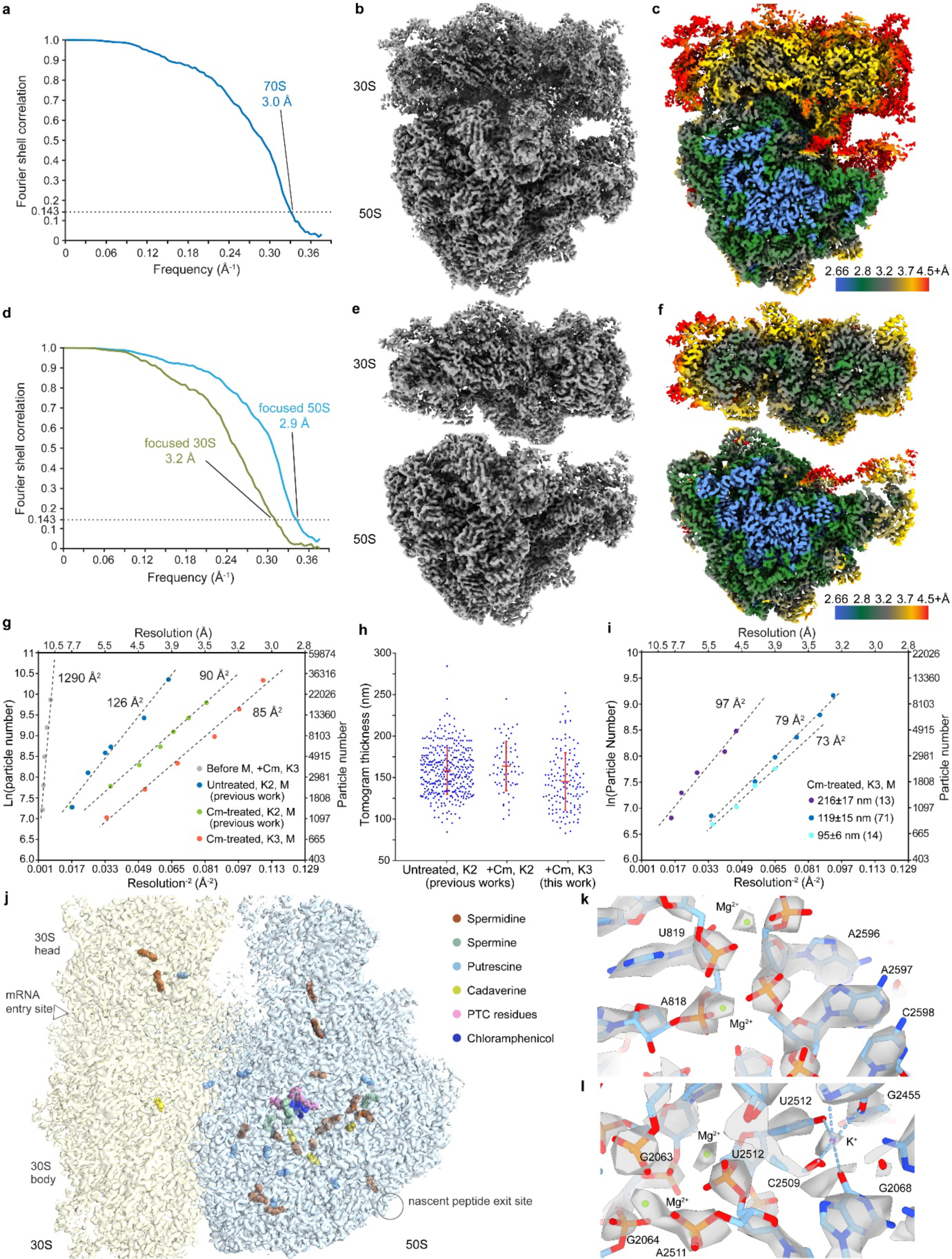
Ribosome sub-tomogram averages in Cm-treated *M. pneumoniae* cells. **a**, Fourier Shell Correlation (FSC) curve for the refined 70S ribosome sub-tomogram average. **b**, The 70S ribosome map. **c**, The 70S map colored by local resolutions. **d**, FSC curves for the small (30S) and large (50S) subunits after focused refinements of the 70S ribosome map. **e**, Maps of the small and large subunits. **f**, Maps of the small and large subunits colored by local resolution. **g**, B factor plots for the dataset processed in the current work in comparison to previously published *M. pneumoniae* ribosome data^15^. **h**, Comparison of cell thickness distribution for the three datasets described in g.. Red lines indicate mean and standard deviation. Cell thickness for the dataset described in this work is 145 ± 35 nm. **i,** B factor plots for selected tomogram subsets of different cell thickness form the current dataset. The cell number, mean and standard deviation values are shown. Analysis of a subset of cells with thickness around 100 nm achieved B factor lower than 80 Å^2^. **j**, Overview of polyamines resolved and modeled in the ribosome structure. **k**, Two Mg^2+^ ions coordinated by 23S rRNA nucleotides^26^. **l**, Region near the PTC showing rRNA nucleotides and ions. Five coordination bonds for the K^+^ ion can be mapped^25^.

**Extended Data Fig. 2.**
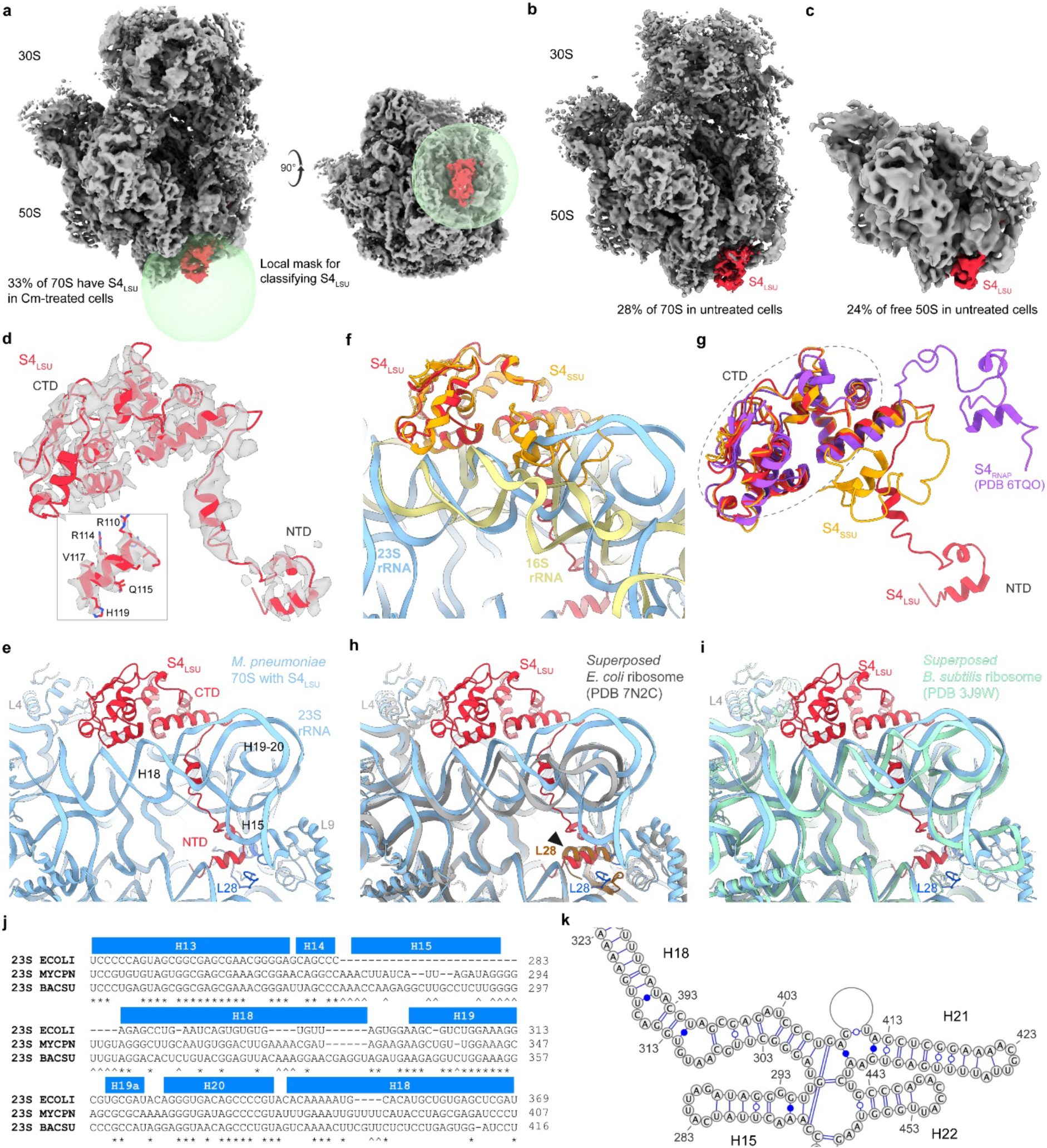
*M. pneumoniae* ribosomes with additional S4_LSU_. **a**, Local mask (green sphere) used for the classification of ribosomes with S4_LSU_ (red) in Cm-treated cells. **b**, In native untreated cells, about 28% of 70S ribosomes have S4_LSU_. **c**, In untreated cells, 24% of free 50S have S4_LSU_. **d**, S4_LSU_ fitted in the ribosome map in Cm-treated cells. Inset: Map and model for a short helix of S4_LSU_ showing the quality of the map allowing robust model building. **e,** Region of the *M. pneumoniae* ribosome structure with S4_LSU_, which contacts 23S rRNA helices 10, 12-16, 18-20 and 52, as well as protein L28. **f**, Comparison of the canonical S4 (S4_SSU_) binding site on the small ribosomal subunit and the S4_LSU_ binding site (aligned on S4). **g,** Comparison of the different conformations of S4 in the RNA polymerase anti-termination complex (purple), the small (yellow) and large ribosomal subunits (red). Models were aligned on S4 CTD, showing that the conformations of the NTD in the 3 models and their relative orientation to the CTD differ. **h,** S4_LSU_ N-terminal helix would clash (black arrowhead) with the ribosomal protein L28’s C-terminal helix of a superposed *E. coli* ribosome which has 13 additional amino acids in the C-terminus compared to *M. pneumoniae* L28. **i,** *B. subtilis* L28 does not have a C-terminal helical extension and could accommodate the superposed S4_LSU_. It also harbors 23S rRNA helix 15 of a similar length to *M. pneumoniae*. **j,** Sequence alignment of 23S rRNA helices 13-20 of *E. coli, B. subtilis* and *M. pneumoniae*, showing the insertion of helix 15 in *M. pneumoniae* and *B. subtilis* compared to *E. coli* (see also panels **e** and **h**). **k,** Secondary structure plot of 23S rRNA helices 14-22 in *M. pneumoniae*.

**Extended Data Fig. 3.**
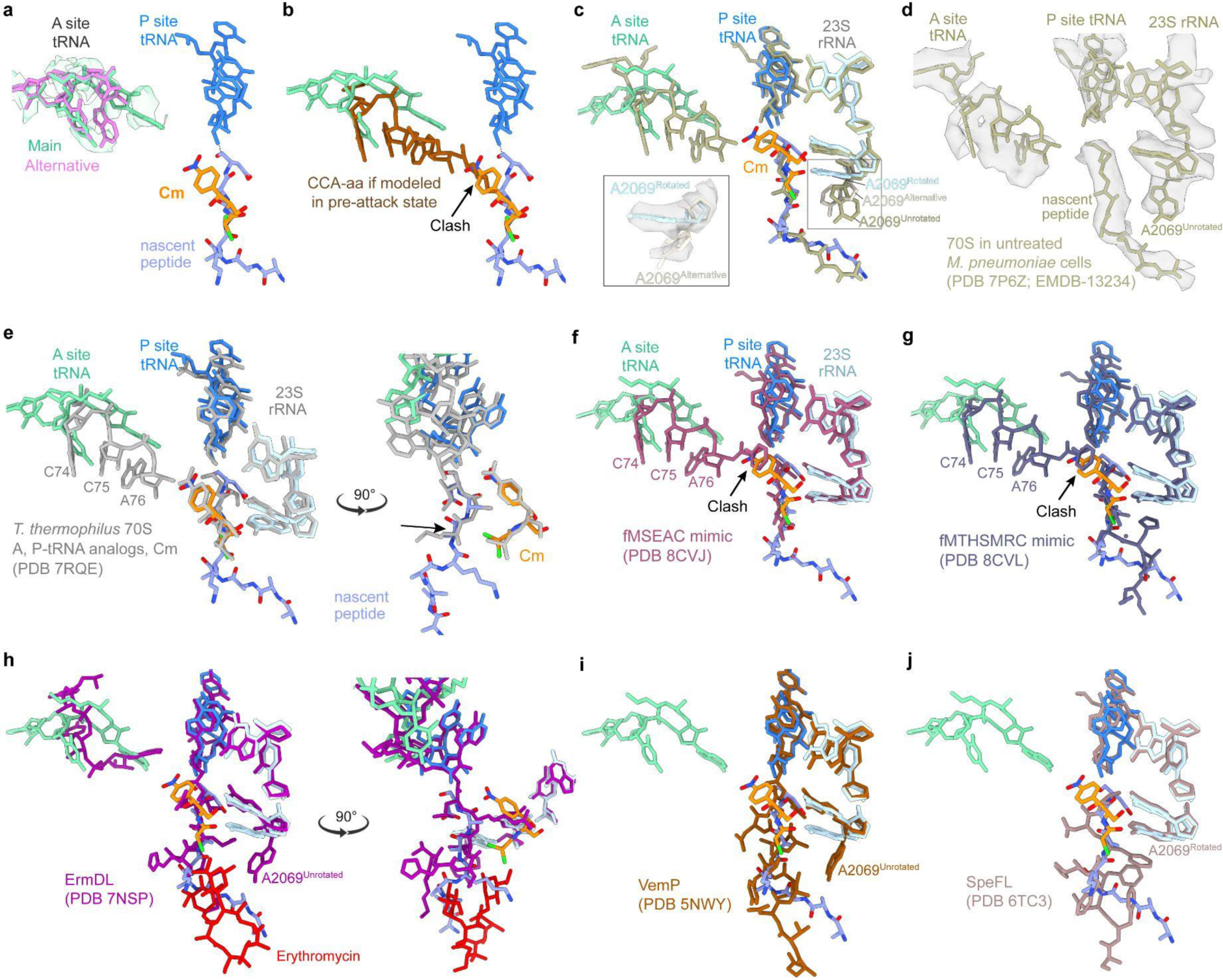
Comparison of the PTC in the presence of Cm across different structures. **a,** Two possible modeling solutions for the CCA-tail (main, green; alternative, magenta) of the aminoacyl-tRNA in the A site. **b,** Clash of the incoming amino acid of A-site tRNA(brown) with Cm (arrow, orange) if modeled in the pre-attack state. **c-d,** Superposition of 70S model from untreated *M. pneumoniae* cells^15^ (yellow-gray) with the Cm-treated 70S model determined in this study. Inset: A2069 in Cm-treated ribosome has an alternative conformation (light gray), similar to the unrotated conformation in the untreated structure. **d,** 70S map and model from untreated *M. pneumoniae*. **e,** Comparison of the Cm-treated 70S model determined in this study with *in vitro* Cm-treated *T. thermophilus* ribosome (grey) with deacylated tRNA analog in A site and non-hydrolyzable peptidyl-tRNA analog that mimics the nascent peptide up to position -2 (black arrow). **f-g,** Comparison of the Cm-treated 70S model determined in this study with *in vitro T. thermophilus* 70S structures with non-hydrolyzable aminoacyl-tRNA^Phe^ and peptidyl-tRNA analogs that mimics the peptide sequences fMSEAC (**f**, purple) and fMTHSMRC (**g**, grey blue). **h-j,** Comparison of the Cm-treated 70S model determined in this study with *in vitro* structures of stalled ribosome-nascent peptide complexes: ErmDL with erythromycin (**h**, dark purple), VemP (**i**, sienna); SpeFL (**j**, tan).

**Extended Data Fig. 4.**
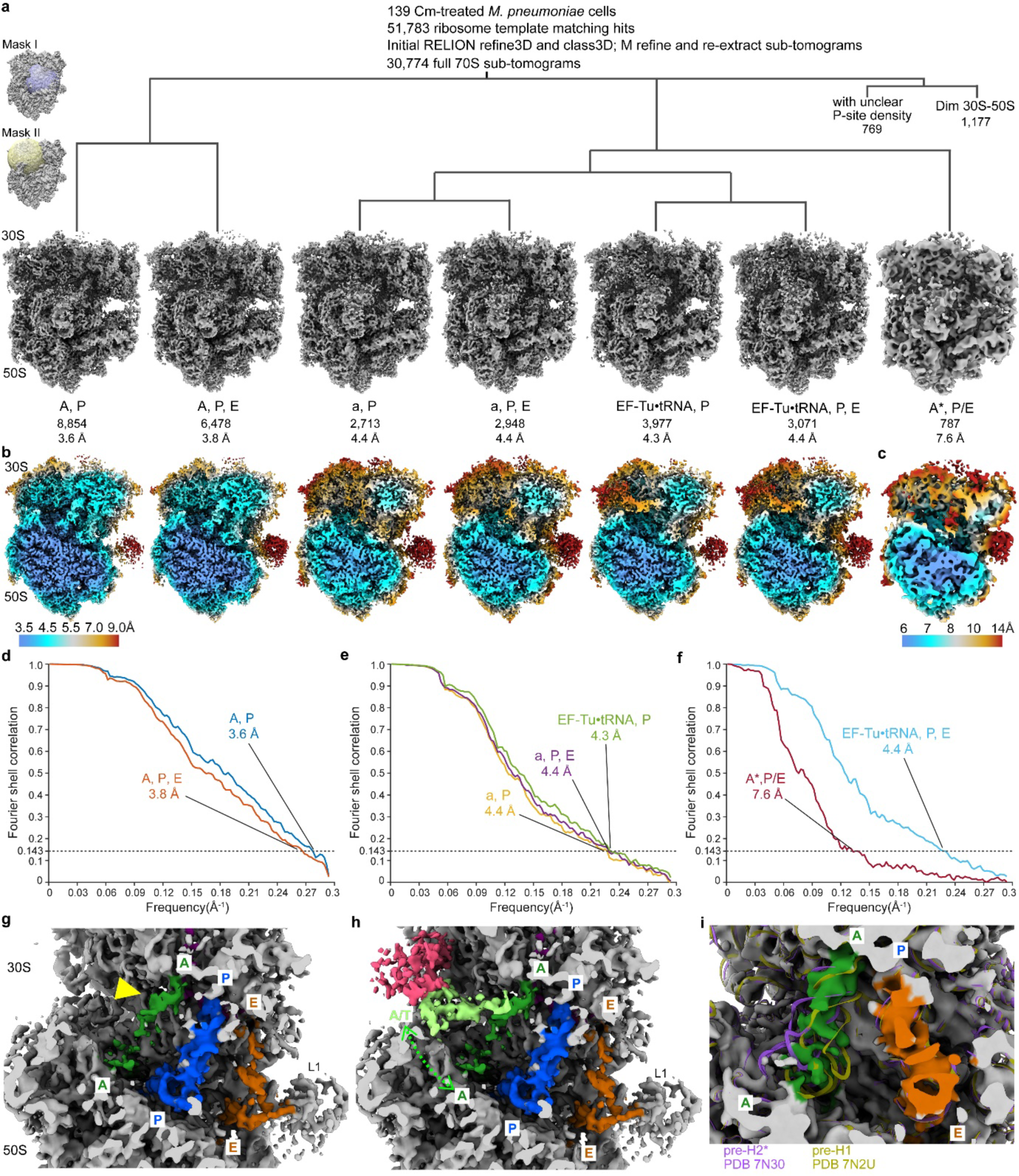
Classification of ribosomes functional states in Cm-treated *M. pneumoniae* cells. **a,** The cryo-ET data processing and sub-tomogram classification procedure. At least three tiers of classifications were performed, including global 70S classification, focused classification on A-P-E tRNA association sites (mask I), and on elongation factor and A/T tRNA binding region (mask II). For each class, the average map, a class name, particle numbers and the global resolution (FSC = 0.143) are provided. Two classes generated at the first tier of global 70S classification (769 and 1,177 particles) resulted in low-resolution ribosome maps with unexplainable densities and are not shown. These are denoted as “unclear” in Extended Data Fig. 5. **b,** Local resolution maps (color coded according to the bar on bottom left) for the six major classes. **c,** Local resolution map (color coded according to the bar on bottom) for the minor class. **d-f,** FSC curves for all classes based on RELION postprocessing. **g-h,** In the “a, P” and “a, P, E” classes, the A-site tRNA’s anticodon end bound to the 30S decoding center is well-resolved (yellow arrowhead), but the body shows blurred density. They possibly represent 70S intermediates with the incoming tRNA sampling between the A/T and the classic A site (as illustrated in **h**). Maps were low-pass filtered to 6 Å for comparison. **i,** The “A*, P/E” class represents intermediates between the pre-translocational hybrid H1 and H2* states^27^. Models are fitted in the map for comparison.

**Extended Data Fig. 5.**
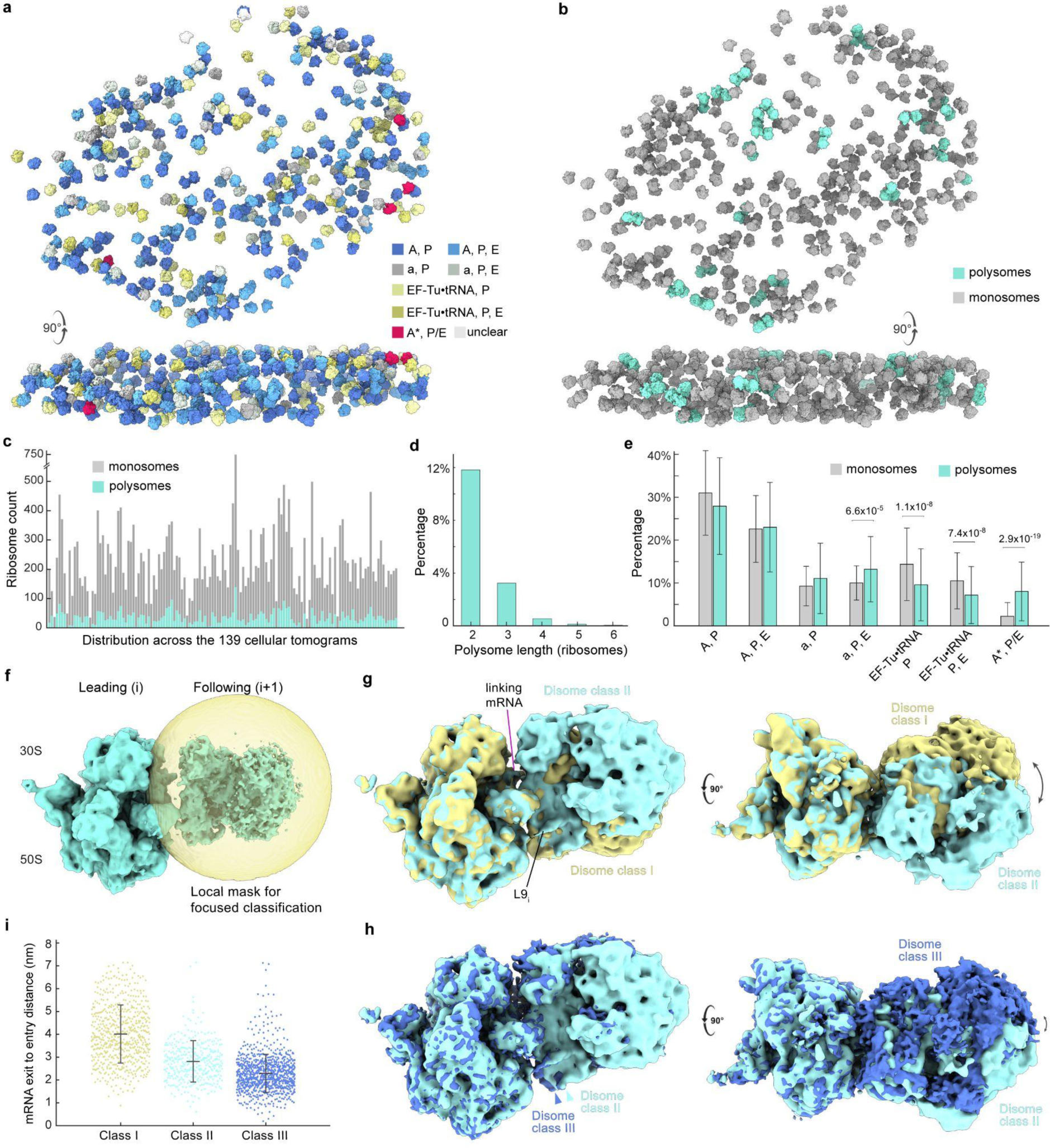
Spatial and structural analysis of polysomes in Cm-treated *M. pneumoniae* cells. **a,** Ribosomes of different functional states classified as described in Extended Data Fig. 4a mapped back into the 3D cellular volume. **b,** The same cell as in panel **a** colored according to the polysome detection results (light green: polysomes, gray: monosomes). **c,** Counts of monosomes and polysomes across the 139 cellular tomograms. **d,** Polysome length distribution. **e,** Distribution of functional states in monosomes and polysomes. Bars represent mean percentages and whiskers represent standard deviations across the 139 cells. False discovery rate-adjusted *p* value (two-sided Wilcoxon rank sum test) is provided. **f,** The local mask used to classify neighboring ribosome pairs (disomes) within the 4,839 annotated polysomes. Aligned on the leading ribosome. **g,** Comparison of disome classes I and II, after alignment on the leading ribosome (left). The major difference comes from the relative rotation of the following ribosome. **h,** Comparison of disome classes II and III. The minor difference comes from the displacement and relative rotation of the following ribosome. **i,** Distribution of the mRNA exit_i_-to-entry_i+1_ distances for the three disome classes. Data for 4,839 annotated polysomes. The mean (line) and standard deviation (whiskers) of distances for the three disome classes are 4.01 ± 1.27 nm, 2.81 ± 0.91 nm, 2.29 ± 0.83 nm.

**Extended Data Fig. 6.**
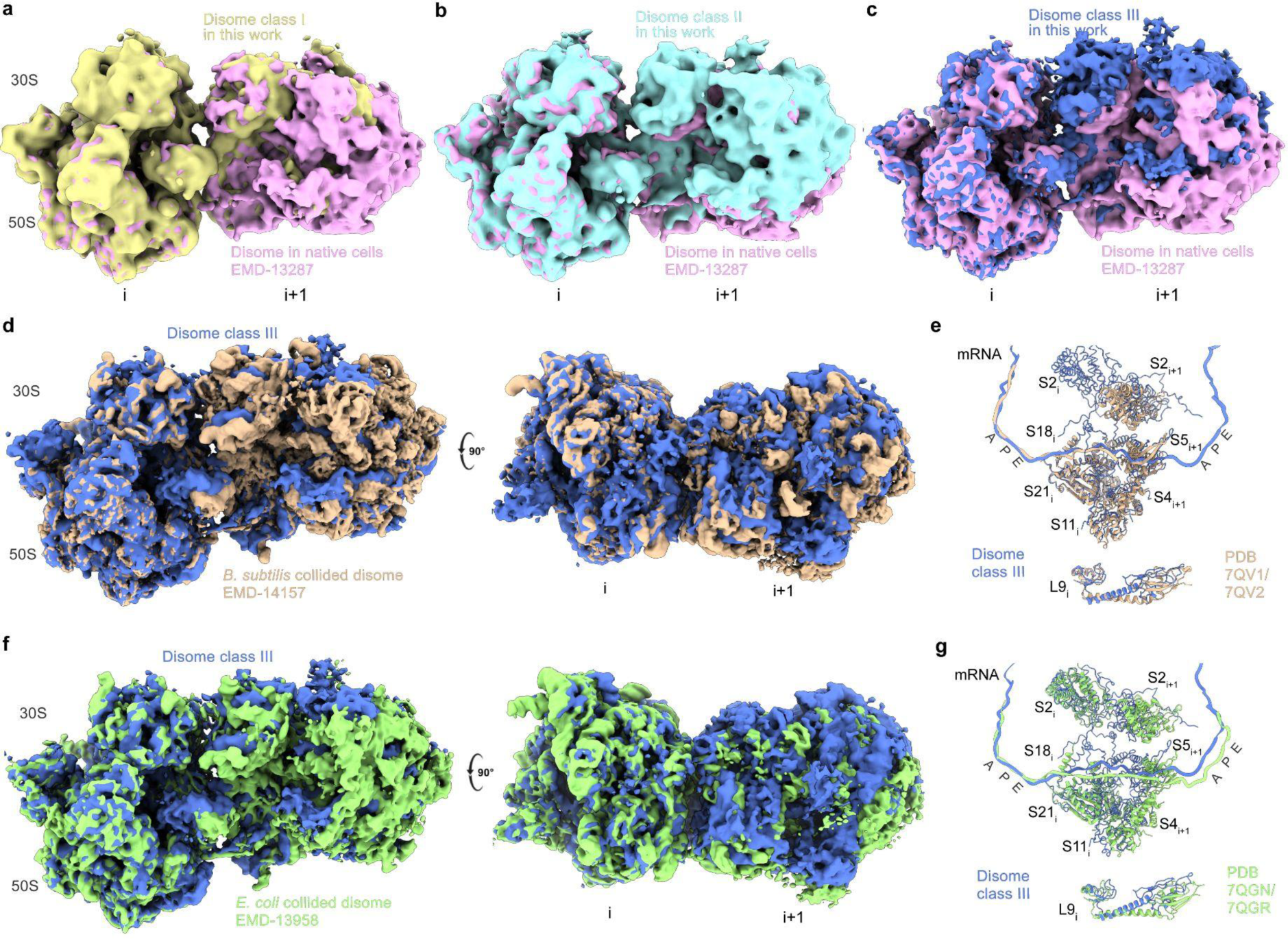
Cm induces collision within polysomes. **a-c,** The Cm-treated disome class I is structurally similar to disomes in native untreated *M. pneumoniae* cells^15^ (**a**), while disome classes II (**b**) and III (**c**) are more compact. The best resolved disome class in untreated cells (pink, EMD-13287) is shown. All maps are aligned on the leading ribosome (left) for comparison. **d-g,** Cm-treated disome class III (blue) resembles the *in vitro* reconstituted collided disomes of *B. subtilis*^29^(**d, e**) and *E. coli*^30^(**f, g**). Aligned on the leading ribosome for comparison. The positioning of mRNA and interface ribosomal proteins largely overlap. There is a slight rotational difference of the L9 protein.

**Extended Data Table 1.** Cryo-ET data collection and structure refinement statistics of high-resolution ribosome averages.

|  | 70S ribosome average<br>in Cm-treated cells<br>EMD-17132<br>PDB 8P7X | 30S small subunit<br>in Cm-treated cells<br>EMD-17133<br>PDB 8P6P | 50S large subunit<br>in Cm-treated cells<br>EMD-17134<br>PDB 8P8B | 70S with a S4<sub>LSU</sub><br>in Cm-treated cells<br>EMD-17135<br>PDB 8P7Y |
| --- | --- | --- | --- | --- |
| **Data collection and processing** |  |  |  |  |
| Magnification | 64,000 |  |  |  |
| Voltage (kV) | 300 |  |  |  |
| Electron exposure (e<sup>-</sup>/Å<sup>2</sup>) | 137 |  |  |  |
| Defocus range (μm) | -1.0 to -3.25 |  |  |  |
| Pixel size (Å) | 1.329 |  |  |  |
| Symmetry imposed | C1 |  |  |  |
| Initial particle images (no.) | 30,774 | 30,774 | 30,774 | 30,774 |
| Final particle images (no.) | 30,774 | 30,774 | 30,774 | 10,151 |
| Map resolution (Å) |  |  |  |  |
| FSC threshold 0.143 | 3.0 | 3.2 | 2.9 | 3.7 |
| Map resolution range (Å) | 2.8 to 6 | 3 to 5 | 2.8 to 4 | 3.2 to 6 |
| Map sharpening B factor (Å<sup>2</sup>) | -1 to +15 | -1 to +15 | -1 to +15 | -1 to +15 |
| **Model refinement** |  |  |  |  |
| Initial model used (PDB code) | 7OOC, 7OOD | 7OOC | 7OOD | 7OOC, 7OOD |
| Model resolution (Å) |  |  |  |  |
| FSC threshold = 0.143 | 3.02 | 3.28 | 3.08 | 3.62 |
| Model resolution (Å) |  |  |  |  |
| FSC threshold = 0.5 | 3.28 | 3.60 | 3.16 | 4.04 |
| Model vs. map correlation<br>coefficient (cc_mask) | 0.73 | 0.74 | 0.80 | 0.73 |
| Model composition |  |  |  |  |
| Non-hydrogen atoms | 151587 | 55468 | 97171 | 153258 |
| Protein residues | 6182 | 2597 | 3578 | 6386 |
| RNA residues | 4755 | 1612 | 3194 | 4755 |
| Ligands/water | 367 | 112 | 266 | 369 |
| B factors (Å<sup>2</sup>) (mean) |  |  |  |  |
| Protein | 70.84 | 82.90 | 62.04 | 71.55 |
| RNA | 73.77 | 88.29 | 67.03 | 73.77 |
| Ligands | 31.27 | 29.99 | 31.44 | 31.27 |
| Water | - | 30.00 | - | - |
| R.m.s. deviations |  |  |  |  |
| Bond lengths (Å) | 0.003 | 0.003 | 0.004 | 0.005 |
| Bond angles (°) | 0.580 | 0.555 | 0.592 | 0.750 |
| Validation |  |  |  |  |
| MolProbity score | 1.93 | 2.03 | 2.20 | 2.10 |
| Clashscore | 8.98 | 10.60 | 7.63 | 13.04 |
| Poor rotamers (%) | 0.83 | 0.58 | 3.75 | 1.00 |
| Ramachandran plot |  |  |  |  |
| Favored (%) | 92.84 | 91.85 | 95.02 | 92.23 |
| Allowed (%) | 6.71 | 7.72 | 4.89 | 7.14 |
| Disallowed (%) | 0.44 | 0.43 | 0.09 | 0.64 |
| Validation (RNA) |  |  |  |  |
| Good sugar pucker (%) | 99.20 | 99.50 | 99.53 | 99.26 |
| Good backbone (%) | 81.42 | 84.40 | 81.70 | 81.49 |

**Extended Data Table 2.** Cryo-ET data collection and structure refinement statistics of ribosome classes.

|  | 70S class<br>"A, P"<br>EMD-17136 | 70S class<br>"A, P, E"<br>EMD-17137 | 70S class<br>"a, P"<br>EMD-17138 | 70S class<br>"a, P, E"<br>EMD-17139 | 70S class<br>"EF-Tu•tRNA<br>, P"<br>EMD-17140 | 70S class<br>"EF-Tu•tRNA<br>, P, E"<br>EMD-17141 | 70S class<br>"A, P/E"<br>EMD-17142 |
| --- | --- | --- | --- | --- | --- | --- | --- |
| **Data collection and processing** |  |  |  |  |  |  |  |
| Magnification | 64,000 |  |  |  |  |  |  |
| Voltage (kV) | 300 |  |  |  |  |  |  |
| Electron exposure (e<sup>-</sup>/Å<sup>2</sup>) | 137 |  |  |  |  |  |  |
| Defocus range (μm) | -1.0 to -3.25 |  |  |  |  |  |  |
| Pixel size (Å) | 1.329 |  |  |  |  |  |  |
| Symmetry imposed | C1 |  |  |  |  |  |  |
| Initial particles (no.) | 30,774 | 30,774 | 30,774 | 30,774 | 30,774 | 30,774 | 30,774 |
| Final particles (no.) | 8,854 | 6,478 | 2,713 | 2,948 | 3,977 | 3,071 | 787 |
| Map resolution (Å)<br>FSC threshold 0.143 | 3.6 | 3.8 | 4.4 | 4.4 | 4.4 | 4.4 | 7.6 |
| Map resolution range (Å) | 3.2 to 4.2 | 3.4 to 4.5 | 4 to 6 | 4 to 6 | 4 to 8 | 4 to 8 | 6 to 10 |
| Map sharpening B factor (Å<sup>2</sup>) | -7 | -5 | -8 | -8 | -1 | -5 | -22 |

**Extended Data Table 3.**
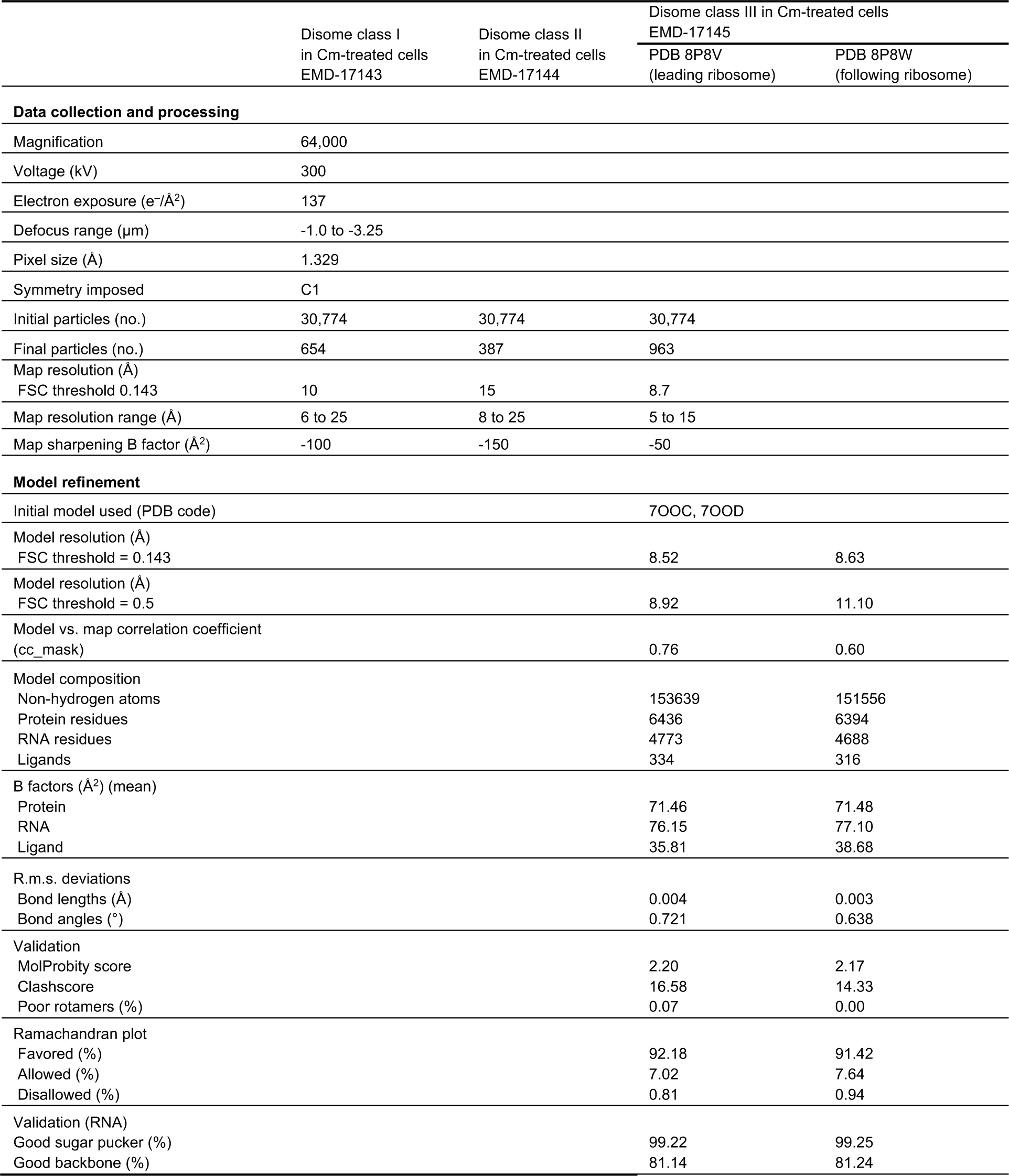
Cryo-ET data collection and structure refinement statistics of di-ribosomes.

|  | Disome class I<br>in Cm-treated cells<br>EMD-17143 | Disome class II<br>in Cm-treated cells<br>EMD-17144 | Disome class III in Cm-treated cells<br>EMD-17145 |  |
| --- | --- | --- | --- | --- |
|  |  |  | PDB 8P8V<br>(leading ribosome) | PDB 8P8W<br>(following ribosome) |
| Data collection and processing |  |  |  |  |
| Magnification | 64,000 |  |  |  |
| Voltage (kV) | 300 |  |  |  |
| Electron exposure (e<sup>-</sup>/Å<sup>2</sup>) | 137 |  |  |  |
| Defocus range (μm) | -1.0 to -3.25 |  |  |  |
| Pixel size (Å) | 1.329 |  |  |  |
| Symmetry imposed | C1 |  |  |  |
| Initial particles (no.) | 30,774 | 30,774 | 30,774 |  |
| Final particles (no.) | 654 | 387 | 963 |  |
| Map resolution (Å) |  |  |  |  |
| FSC threshold 0.143 | 10 | 15 | 8.7 |  |
| Map resolution range (Å) | 6 to 25 | 8 to 25 | 5 to 15 |  |
| Map sharpening B factor (Å<sup>2</sup>) | -100 | -150 | -50 |  |
| Model refinement |  |  |  |  |
| Initial model used (PDB code) |  |  | 7OOC, 7OOD |  |
| Model resolution (Å) |  |  |  |  |
| FSC threshold = 0.143 |  |  | 8.52 | 8.63 |
| Model resolution (Å) |  |  |  |  |
| FSC threshold = 0.5 |  |  | 8.92 | 11.10 |
| Model vs. map correlation coefficient<br>(cc_mask) |  |  | 0.76 | 0.60 |
| Model composition |  |  |  |  |
| Non-hydrogen atoms |  |  | 153639 | 151556 |
| Protein residues |  |  | 6436 | 6394 |
| RNA residues |  |  | 4773 | 4688 |
| Ligands |  |  | 334 | 316 |
| B factors (Å<sup>2</sup>) (mean) |  |  |  |  |
| Protein |  |  | 71.46 | 71.48 |
| RNA |  |  | 76.15 | 77.10 |
| Ligand |  |  | 35.81 | 38.68 |
| R.m.s. deviations |  |  |  |  |
| Bond lengths (Å) |  |  | 0.004 | 0.003 |
| Bond angles (°) |  |  | 0.721 | 0.638 |
| Validation |  |  |  |  |
| MolProbity score |  |  | 2.20 | 2.17 |
| Clashscore |  |  | 16.58 | 14.33 |
| Poor rotamers (%) |  |  | 0.07 | 0.00 |
| Ramachandran plot |  |  |  |  |
| Favored (%) |  |  | 92.18 | 91.42 |
| Allowed (%) |  |  | 7.02 | 7.64 |
| Disallowed (%) |  |  | 0.81 | 0.94 |
| Validation (RNA) |  |  |  |  |
| Good sugar pucker (%) |  |  | 99.22 | 99.25 |
| Good backbone (%) |  |  | 81.14 | 81.24 |

**Extended Data Table 4.** Modeled rRNA modifications and polyamines in *M. pneumoniae*.

| Positions in *M. pneumoniae* | rRNA modification or polyamine types | Corresponding positions in *E. coli* | Corresponding positions in *T. thermophilus* |
| --- | --- | --- | --- |
| 23S rRNA |  |  |  |
| m<sup>1</sup>G783 | 1-methylguanosine | G750, unmodified | G748, unmodified |
| Gm2259 | 2'-O-methylguanosine | Gm2251 | Gm2263 |
| m<sup>2</sup>A2511 | 2-methyladenosine | m<sup>2</sup>A2503 | m<sup>2</sup>A2515 |
| 16S rRNA |  |  |  |
| m<sup>7</sup>G525 | 7-methylguanosine | m<sup>7</sup>G527 | m<sup>7</sup>G527 |
| m<sup>4</sup>C1377 | N<sup>4</sup>-methylcytosine | m<sup>4</sup>Cm1402 | m<sup>4</sup>Cm1402 |
| m<sup>5</sup>C1375 | 5-methylcytosine | C1400, unmodified | m<sup>5</sup>C1400 |
| m<sup>6</sup><sub>2</sub>A1493 | N<sup>6</sup>,N<sup>6</sup>-dimethyladenosine | m<sup>6</sup><sub>2</sub>A1518 | m<sup>6</sup><sub>2</sub>A1518 |
| m<sup>6</sup><sub>2</sub>A1494 | N<sup>6</sup>,N<sup>6</sup>-dimethyladenosine | m<sup>6</sup><sub>2</sub>A1519 | m<sup>6</sup><sub>2</sub>A1519 |
| Polyamines in 50S subunit (appending to chain 3) |  | Compared to *E. coli* ribosome (PDB 7N1P) |  |
| 4007 | Putrescine | Unique |  |
| 4008 | Putrescine | Same orientation |  |
| 4009 | Putrescine | Same orientation |  |
| 4010 | Putrescine | Spermidine in 7N1P, different orientation |  |
| 4011 | Putrescine | Unique |  |
| 4012 | Spermine | Unique |  |
| 4013 | Spermidine | Putrescine in 7N1P |  |
| 4015 | Spermidine | Putrescine in 7N1P |  |
| 4018 | Spermidine | Unique |  |
| 4019 | Putrescine | Unique |  |
| 4020 | Spermidine | Unique |  |
| 4021 | Spermidine | Unique |  |
| 4022 | Spermidine | Unique |  |
| 4023 | Spermidine | Unique |  |
| 4025 | Spermidine | Unique - S58 / *E.coli* N59 chain p (RL20) |  |
| 4029 | Spermidine | Unique - Gln 157 chain b (RL3) |  |
| 4030 | Spermidine | Unique - stacking against A518 (23S) |  |
| 4031 | Spermidine | Unique - stacking against Y139 (RL3) |  |
| 4032 | Spermidine | Unique |  |
| 4033 | Spermine | Unique |  |
| 4034 | Putrescine | Different orientation |  |
| 4035 | Cadaverine | Replaces K111 side chain of L13 protein (D114 in *M. Pneumoniae*) |  |
| 4036 | Cadaverine | Spermidine in 7N1P |  |
| 4038 | Spermidine | Different orientation |  |
| 4039 | Spermidine | Different orientation |  |
| 4040 | Cadaverine | Unique - K11 chain 0 (RL34) |  |
| 4041 | Spermidine | Unique |  |
| 4042 | Spermine | Putrescine in 7N1P |  |
| 4043 | Spermidine | Unique |  |
| 4044 | Spermine | Unique |  |
| Polyamines in 30S subunit (appending to chain 5) |  |  |  |
| 1601 | Spermidine | Unique |  |
| 1602 | Putrescine | Same orientation |  |
| 1603 | Cadaverine | Unique |  |
| 1604 | Spermidine | Unique |  |
| 1605 | Putrescine | Unique |  |

